# Influence of camera type, height and image enhancement on photogrammetry success in turbid marine environments

**DOI:** 10.1101/2024.09.15.613158

**Authors:** K.L. Savill, I. Parnum, J. McIlwain, D. Belton

## Abstract

Over the last decade, Structure-from-Motion (SfM) photogrammetry has successfully been used to survey marine benthic environments, including artificial reefs, shipwrecks, and coral reefs, for a wide range of applications. The method is likely to become one of the most common tools for surveying marine benthic environments. However, SfM photogrammetry has been developed in clear water environments, and its suitability in turbid, benthic environments is less certain. Turbid coral reefs are example of an important marine benthic environment, making up 12% of coral reefs globally. Corals in these environments have a tolerance for low-light and high sediment conditions. Such attributes mean these reefs may be important refuges from extreme light and temperatures. Therefore, assessment and optimisation of the photogrammetry methodology in these environments is needed. This study investigates the performance of SfM photogrammetry in turbid environments, by comparing two camera types, settings (automatic vs manual derived from local conditions), the height of image acquisition above the seafloor, and post-processing image enhancement. Three dimensional (3D) SfM photogrammetry meshes of an artificial reef structure using two cameras, an action camera and a compact camera, were compared with its known dimensions detailed in an engineering diagram. According to surface area calculations, the compact camera provided a better 3D mesh than the action camera, with surface area calculations providing an accuracy of 98.2% against the engineering model, compared to 93.2% for the action camera. Images taken at a height of 1 m above the seafloor provided 3D meshes that were more accurate than those using images taken at 2 m above the seafloor. Two image enhancement techniques, histogram equalisation and contrast limited adaptive histogram equalisation (CLAHE), were then applied to assess if this improved the SfM photogrammetry mesh. The 3D mesh from images using the action camera that had a histogram equalisation enhancement provided the most comparable surface area measurement to the engineering diagram, with 100.6% accuracy, indicating our mesh had accounted for growth of benthic organisms on the structure since its installation. In contrast, raw (not enhanced) images had most comparable surface area measurement (98.2% accuracy) using the compact camera. However, the higher apparent accuracy of surface area measurements with the action camera following image enhancement may also be an artefact of inaccurate visual representations from the 3D mesh. Given the comparable accuracy of both approaches, we suggest SfM photogrammetry in turbid benthic environments uses cameras with a larger sensor sizes and customisable settings. This will result in the most accurate 3D meshes from raw imagery, particularly with images taken at a close distance (e.g., ≤ 1 m above the seafloor) and at high intervals (0.5 sec) with percentage overlap (>70%) among images. As the artificial reef in this study was in shallow water (3-4m), lights and/or strobes should be taken into consideration in deeper turbid waters but can also cause problems such as backscatter. Lastly, image enhancement can provide a means to improve image quality, and overall 3D mesh accuracy, when raw image quality and choice of cameras settings were poor.

## 1. INTRODUCTION

With rapid advances in digital technology, the use of 3D SfM photogrammetry has been growing in marine systems over the last decade and has become a common method to survey marine benthic environments, such as coral reefs. The benefits of SfM photogrammetry are that it is a non-invasive, relatively low-cost method that creates high-resolution models which can be repeated to monitor change over time and explore many other ecological interactions (Lange et al., 2020; Roach et al. 2021). This method has been developed to quantify 3D characteristics of coral reefs (e.g., Burns et al., 2019), allowing assessments of reef health (e.g., Gibson, 2021), response to disturbance events (e.g., Longo et al., 2020; Fukunaga et al., 2022) and general monitoring of background changes (e.g., Lange et al., 2020; Rossi et al., 2020). The main advantage is the ability to extract many different types exploring community dynamics on the reef, such as changes in percentage cover of different group using point-intercept methods, or tracking demographic (growth, survival) changes in individuals responding to physical or biological interactions.

The applications to documenting community changes range from monitoring coral reefs to tracking community development on artificial structures (Abadie et al., 2018; Rofallski et al., 2020; Fogg and McDougall 2023). Yet, previous studies using this technique have mostly taken place in clear water environments, with few applications in turbid benthic environments.

SfM photogrammetry has been in the terrestrial environment since the late 1990’s, but underwater environments pose challenges for photography, such as low light, low contrast, loss of colour and noise (Arnold, 2022), and specialised camera equipment to withstand saltwater and high pressures. These challenges are even greater in turbid water environments which have significantly less light, poor visibility and a significant amount of particulate matter in the water column (IADC, n.d.) compared to clear water environments. Yet, turbid water environments represent ∼8 to 12% of the total area of the global continental shelf (Shi and Wang, 2010) but are relatively understudied, in part due to methodological challenges with surveying these systems. Thus, there is an important need to trail and share methods to improve photography in low light environments.

In general, there are three stages to improve image acquisition and model accuracy in SfM photogrammetry; equipment choice (camera and additional equipment); image acquisition (e.g., settings, image replication, distance from substrata) and post-processing to improve image quality (e.g., image enhancement). This study investigates methods of image collection in turbid benthic environments and post-processing to help improve the performance of SfM photogrammetry, particularly the accuracy of 3D reconstructions. This was done by re-surveying an artificial reef of known dimensions in turbid conditions with different camera types, camera settings, image numbers, height from the substrate. The effect of post-processing image enhancement on feature alignment and model accuracy were also investigated. Model accuracy was assessed by comparing reconstructions of the artificial reefs in field surveys with the engineering diagram of the structures.

## 2. METHODS

### 2.1 Study area

This study was conducted at the Coogee Maritime Trail at Coogee Beach, Perth, Western Australia (Figure 1). This site was chosen as it is relatively exposed and often experiences low visibility and turbid conditions due to prevailing winds and local wind-driven waves. In addition, the study site contains prefabricated structures (artificial reefs) with known dimensions, such as the ‘Apollo’ cluster structures (Figure 1d), which provided a way to determine the accuracy of the SfM photogrammetry methodology. These first structures of the maritime trail were installed in 2016, while the ‘Apollo’ cluster surveyed was installed in 2019, with 3 years of external growth at the time of survey.

**Figure 1.**
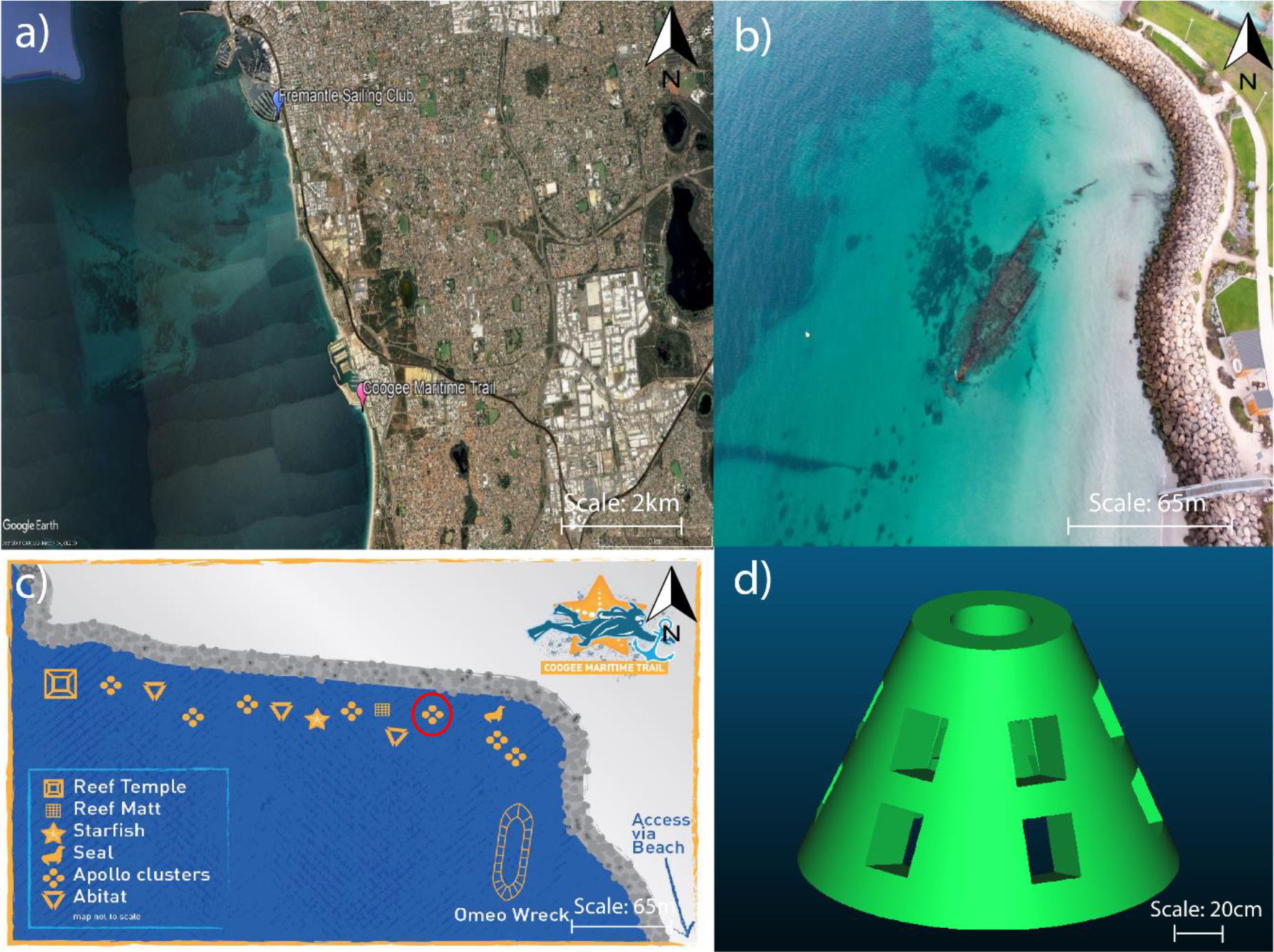
Coogee Maritime Trail: a. location on the Western Australian Coastline south of Perth city (Google Earth Pro); b. aerial photograph (Seniorocity, 2020); c. map of the artificial reef structures (Tomlinson, 2022), with red circle indicating surveyed structure; d. 3D engineering diagram of artificial reef structure (Subcon Engineering).

### 2.2 Cameras

Two types of cameras were used to capture images of the artificial reef structure: an action camera (GoPro Hero 8) and compact camera (Canon G7X Mark II).

**Table 1.** Specifications of cameras used in this study.

| Specification | Camera |  |
| --- | --- | --- |
|  | GoPro Hero 8 | Canon G7X Mark II |
| Cost (as of Aug 2024) | \$398 AUD | \$1,199 AUD |
| Sensor resolution (Megapixels) | 12 | 20 |
| Sensor size (mm) | 6.17 x 4.55 | 13.2 x 8.8 |
| Image size (pixels) | 4000x3000 | 5472x3648 |
| Focal length (35mm equivalent) | 19-39 mm (Linear mode) | 24-100 mm |

### 2.3 Data collection and processing

Turbidity of the study site was taken into consideration when selecting the camera settings. Turbidity was quantified as suspended sediment (NTU) during the survey period (day/time/hrs) with XX instrument/sensor. The action camera set to linear mode and the compact camera was set to a focal length of 24 mm, with images of the artificial structure taken with both cameras (one after the other) continuously at a rate of ∼1 m/s, firstly at a height of 1m and then 2m above the seafloor (Figure 2). The 3D mesh of the artificial reef was created using AgiSoft Metashape (Version 1.7.1), using the workflow from Lange and Perry (2020). In AgiSoft Metashape, the camera calibration and optimization resolves the optical characteristics of the camera lens using the image metadata and Brown’s distortion model (Burns et al., 2015). Images for each of the 18 reconstructions (camera type x height above seafloor x image enhancement) were aligned and a 3D mesh was created. These were then overlaid on each other and clipped to exclude the surrounding seafloor, allowing for surface area calculations of the artificial reef structure only (Figure 3).

**Figure 2.**
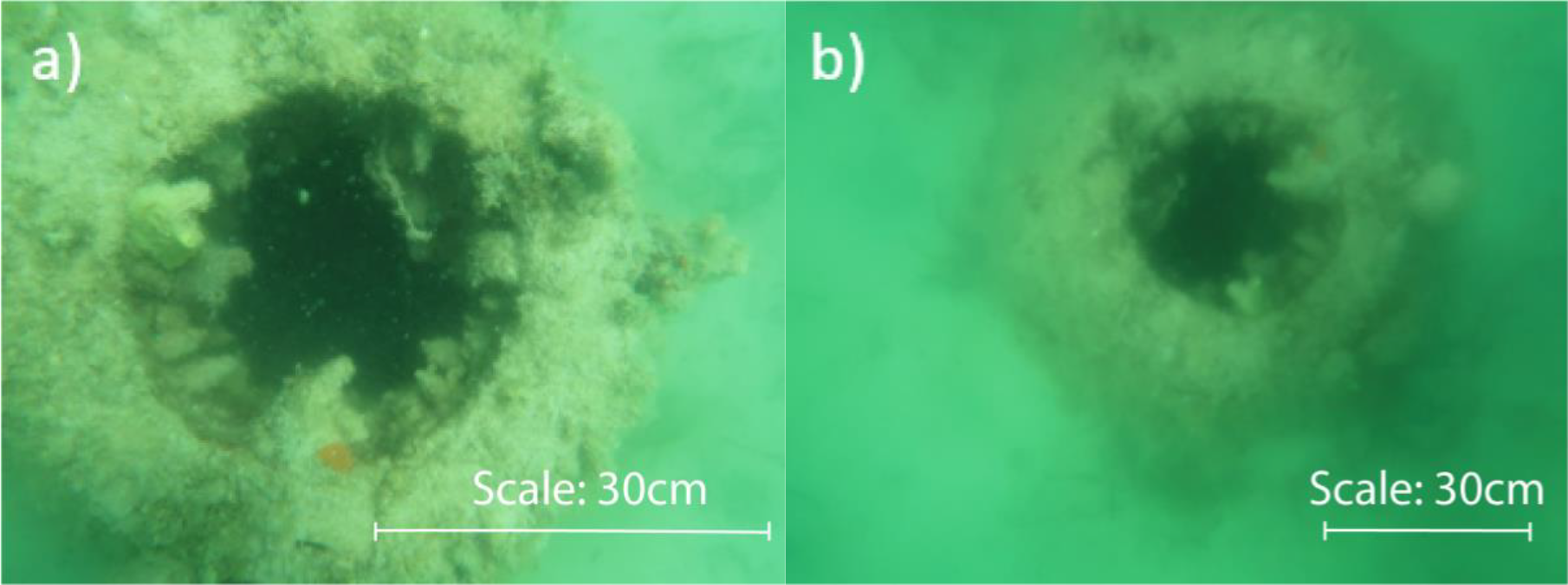
Example of images taken with compact camera at **a**. 1m above the seafloor, and **b**. 2m above the seafloor, of the artificial reef structure.

**Figure 3.**
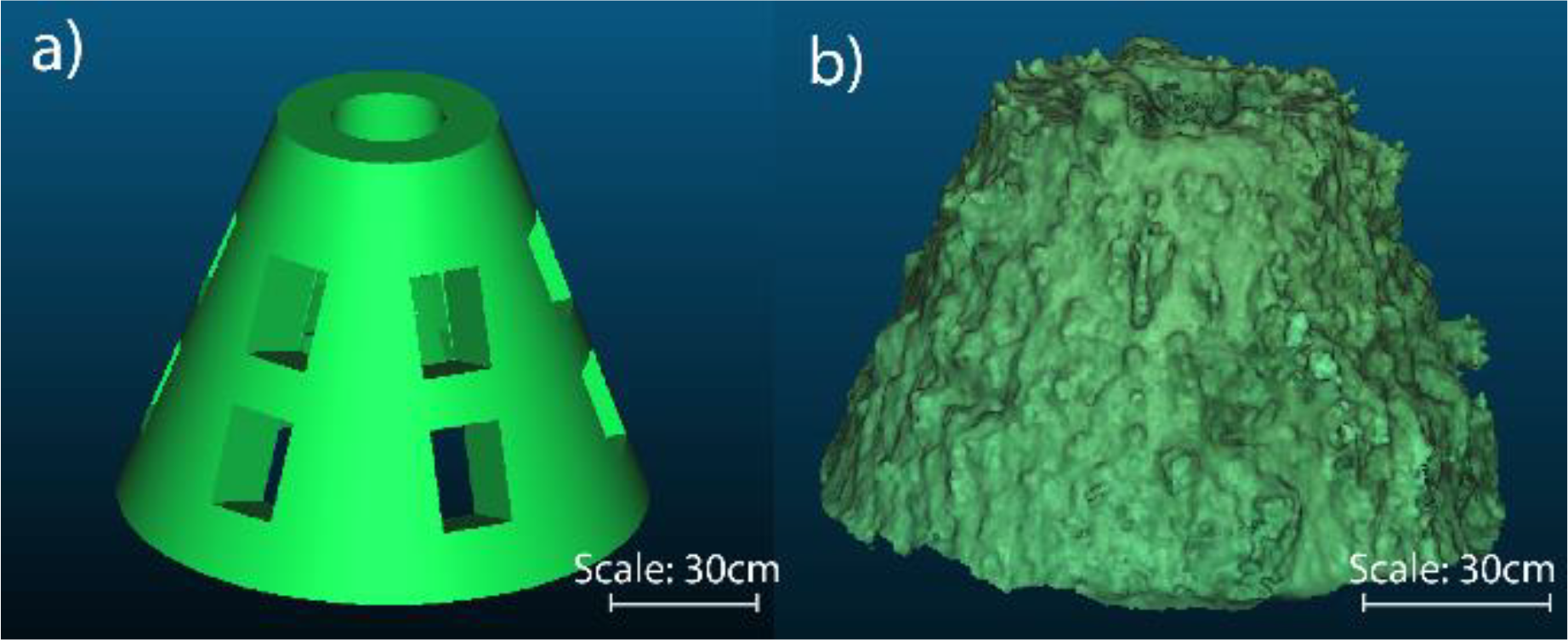
**a**. Engineering diagram; **b**. 3D mesh from compact camera from images taken at both 1m and 2m.

### 2.5 Data analyses

To determine the surface area of the point cloud measurements, for the engineering model and the 3D meshes from in-situ imagery, each mesh created was measured within the software CloudCompare (Version 2.12). The percentage agreement (surface area of 3D mesh compared to the surface area of the engineering model) was used as a proxy for accuracy and a regression analysis was carried out in MATLAB (R2021a) using the built in “regress” function and the Root-Mean-Squared-Error (RMSE) was used to assess regression model performance. Then, to determine the accuracy of the surface are calculation of the meshes against the number of images used to create it and the distance above the seafloor they were taken at (i.e., 1m or 2m), a multi-linear regression analysis was conducted. The resulting 3D models were then compared to see how well the surface area agreed with the engineering diagram.

## 3. RESULTS

### 3.1 Environment and turbidity

During data collection, Coogee Maritime Trail had a range of 9-16 NTU and an average of 11.5 NTU, although the in-situ visibility appeared higher, with a visual line of site limited to 1.5-2m (Figure 2).

### 3.2 Surface area comparisons

Of the 3D meshes created of the artificial reef structure created from the field surveys, both the action camera and compact camera 3D meshes using images from both 1m and 2m above the seafloor produced a surface area calculation that was the closest to that of the engineering model (Table 2).

**Table 2.**
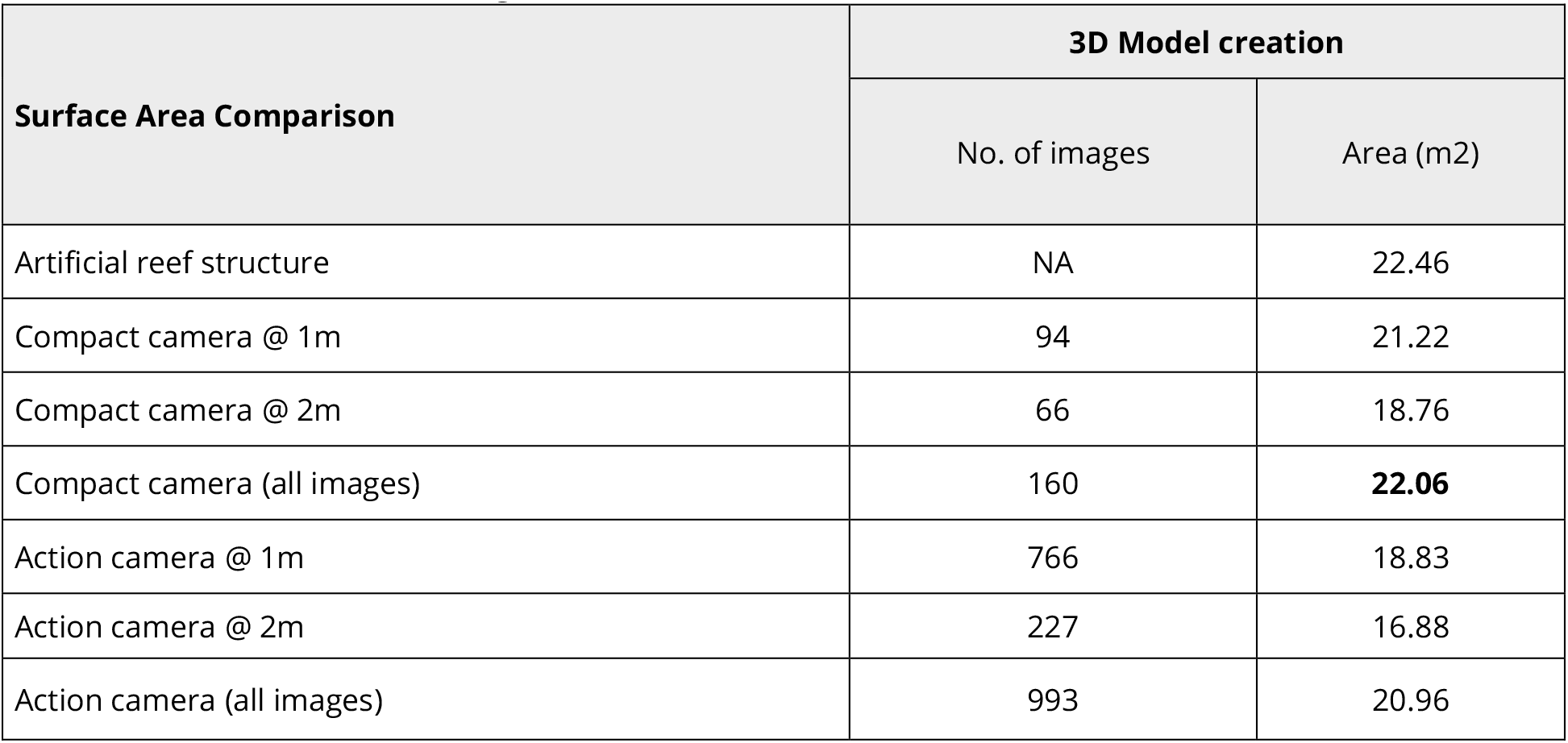
Number of images and surface area measurements (m_2_) of the artificial reef structure from the compact camera and action camera created using images from 1m, 2m and combined 1m & 2m above the structure. Bold text indicates the highest surface area measurement calculation.

The 3D mesh created using images from the action camera at 1 m above the seafloor with a histogram equalisation applied to the images provided the most accurate surface area measurement to the engineering model, at 100.6% (Table 4.2). The 3D mesh created using raw images taken at both 1m and 2m above the seafloor using the compact camera, provided the second most accurate surface area measurement to the engineering model, with 98.2% (Table 4.2).

**Table 4.2.** Surface area comparison of Apollo structure engineering diagram against 3D models created from action camera and compact camera using original, histogram equalisation and CLAHE images. Bolded values indicating the most agreeable surface area measurement.

| Surface Area Comparison | 3D Model creation |  |  |  |
| --- | --- | --- | --- | --- |
|  | No. of images | Original | Histogram | CLAHE |
| Artificial reef structure | NA | **22.46** | NA | NA |
| Compact camera @ 1m | 94 | 21.22 | 47.58 | 15.38 |
| Compact camera @ 2m | 66 | 18.76 | 28.4 | 19.62 |
| Compact camera (all images) | 160 | **22.06** | 16.52 | 19.06 |
| Action camera @ 1m | 766 | 18.83 | **22.6** | 42.8 |
| Action camera @ 2m | 227 | 16.88 | 7.46 | 52.06 |
| Action camera (all images) | 993 | 20.96 | 8.11 | **24.09** |

### 3.5 Regression analysis

The predicted accuracy of 3D models from linear and multilinear regression of the number of images at different heights above the seafloor. Accuracy of the surface area calculations from the 3D meshes could be well approximated by using both regressions, however, the multilinear regression found that images taken at a height of 1 m above the seafloor had more bearing on the accuracy of the 3D model, than images taken at 2 m, therefore the closer the image was taken to above the seafloor, the more accurate the 3D mesh surface area calculations were likely to be. Linear and logarithmic regression was used to look at the relationship between the total number of images and measurement accuracy, and although the RMSE for a linear and log functions were both 5%, the best visual fit was using a log relationship (Figure 5).

**Figure 5.**
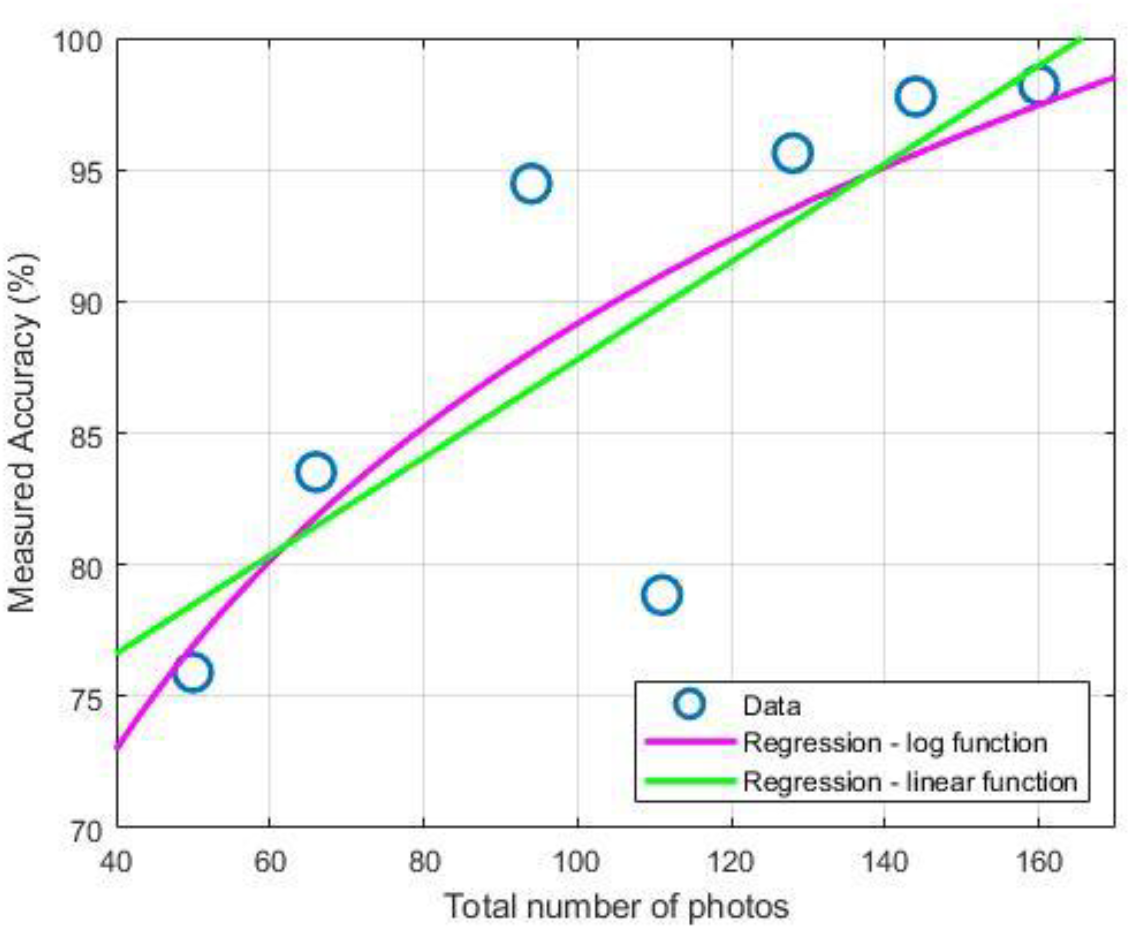
3D mesh generated using different numbers of images (blue data points) with a linear regression (green line) of the measured accuracy against total number of images, and a line of best fit using a log relationship (pink line).

## 4. DISCUSSION

While we were able to generate 3D meshes for all scenarios we trialled, it was found that the compact camera, outperformed the action camera, producing the most accurate surface area calculation, closest to that of the engineering model. Action cameras have benefits such as being *significantly cheaper than a compact camera, easy to use, relatively automatic, extremely compact and relatively robust* and have been used in various previous studies on coral reefs (Cresswell et al., 2020) and other places. However here we show that using a compact camera is worthwhile depending on the research question. For instance, if measuring objects from the 3D meshes, it is likely that more accuracy can be obtained from the compact camera, as shown in this study from the surface area calculations of the 3D meshes against the artificial reef structure surface area from the engineering diagram. We suggest the reasons for the compact camera performing better include the camera’s large sensor and image size, as well as the custom settings used as outline in the methos, makes this camera ideal for turbid benthic environments, as it has larger photosites, a light sensitive element on the sensor (McCarty, n.d.), allowing for high quality images in low light situations (Maio, 2020).

From a review of relevant literature from open-access journals from 2015 to 2022, it was identified from 43 studies of photogrammetry, only eight used a camera with comparable sensor and image size to the compact camera in this study. The most commonly used cameras were a version of the action camera, which were used in 17 of 43 studies. Of these studies, most were conducted in clear water environments, where conditions are less challenging for SfM photogrammetry than in turbid benthic environments.

Previous studies using compact cameras have used custom settings to suit the environment they are surveying in (e.g., Burns et al., 2015; Burns et al., 2017; Fukunaga et al., 2019; Suka et al., 2020; Roach et al., 2021; Pascoe et al., 2021). This would suggest that if a camera has customisable settings (i.e., can manually adjust to suit different conditions), it will provide higher quality imagery than that of the automated settings, such is the case for most action cameras.

This study found evidence that images taken at closer height above the seafloor in a turbid benthic environment, helped improve the accuracy of surface area calculations from the 3D meshes produced. However, in general it was found that more images from either height increased model accuracy to a point of 120 images, i.e., the rate of increase of model accuracy eventually stabilises from 120 images. It is possible, though, that the mixture of heights from the seafloor improved the estimation of the camera’s internal orientation parameters as part of the SfM photogrammetry bundle adjustment, but this was not conclusive.

When image quality is less than optimal, due to available cameras or limited camera settings, image enhancement techniques can be used to enhance image quality and improve the accuracy of surface area measurements from 3D meshes. The results on image enhancement techniques used to improve accuracy of the surface area calculations from the 3D meshes in this study reflect those of previous studies (e.g.,Li et al., 2015; Xu et al., 2016; Alasal et al., 2018; Qiao et al., 2019; Xu et al., 2020), however, as our image enhancement only improved the accuracy of the 3D mesh surface area calculations for the action camera, it appears that the CLAHE method used in this study may require more investigation to optimise its use in turbid underwater environments. The simplistic histogram equalisation improves the images taken closest to the seafloor and is suited improving image quality and subsequent 3D meshes for turbid water environments.

### 4.6 Uncertainty in findings

The 3D meshes created of the artificial reef structure at Coogee Maritime Trail showed that the compact camera produced the most agreeable surface area calculation of the artificial reef, producing the highest surface area measurement of all the 3D meshes. The 3D mesh slightly underestimated the surface area, which is the opposite of what we might expect, given the surface area of the artificial reef structure had growth from algae, tunicates, and other organisms. The artificial reef was deployed in 2019 and growth of plant and animal communities was evident when comparing the reconstruction from the engineering diagram and that from the 3D meshes from our field surveys (Figure 3). Conversely, the additional area and apparent growth of benthic organisms on the artificial reef over 3 years may be visual noise (e.g., Figure S1) that could be attributed to the histogram stretching of the images causing the SfM photogrammetry program to create pseudo-features. In contrast, the 3D meshes which were incomplete (i.e., had holes; Figure S2) subsequently had a lower surface area measurement. The ability of SfM photogrammetry 3D meshes to accurately represent structures of known area, and additional areas or volumes attributable to growth of marine organisms requires further investigation. Indeed, the ultimate goal of the 3D meshes of coral communities on turbid reefs it to produce reconstructions and images of sufficient quality to distinguish areas among different benthic organisms and to quantify changes in through time following periods of disturbance and recovery. The capacity to distinguish benthic groups with the required image quality is a particular challenge on turbid reefs, due to lack of light availability and suspended particulate in the water column, which is why a methodology for SfM photogrammetry in these environments is so crucial to allow for accurate analysis of growth on these reefs.

## SUPPLEMENTARY MATERIAL

### Data S4.1. Histogram Equalisation

Histogram equalisation of images was executed in MATLAB prior to alignment. Relevant code is available on figshare [https://doi.org/10.6084/m9.figshare.20509122.v1]

### Data S4.2. CLAHE

CLAHE of images was executed in Juptyer Notebook prior to alignment. Relevant code is available on figshare [https://doi.org/10.6084/m9.figshare.20509122.v1]

**Figure S1.**
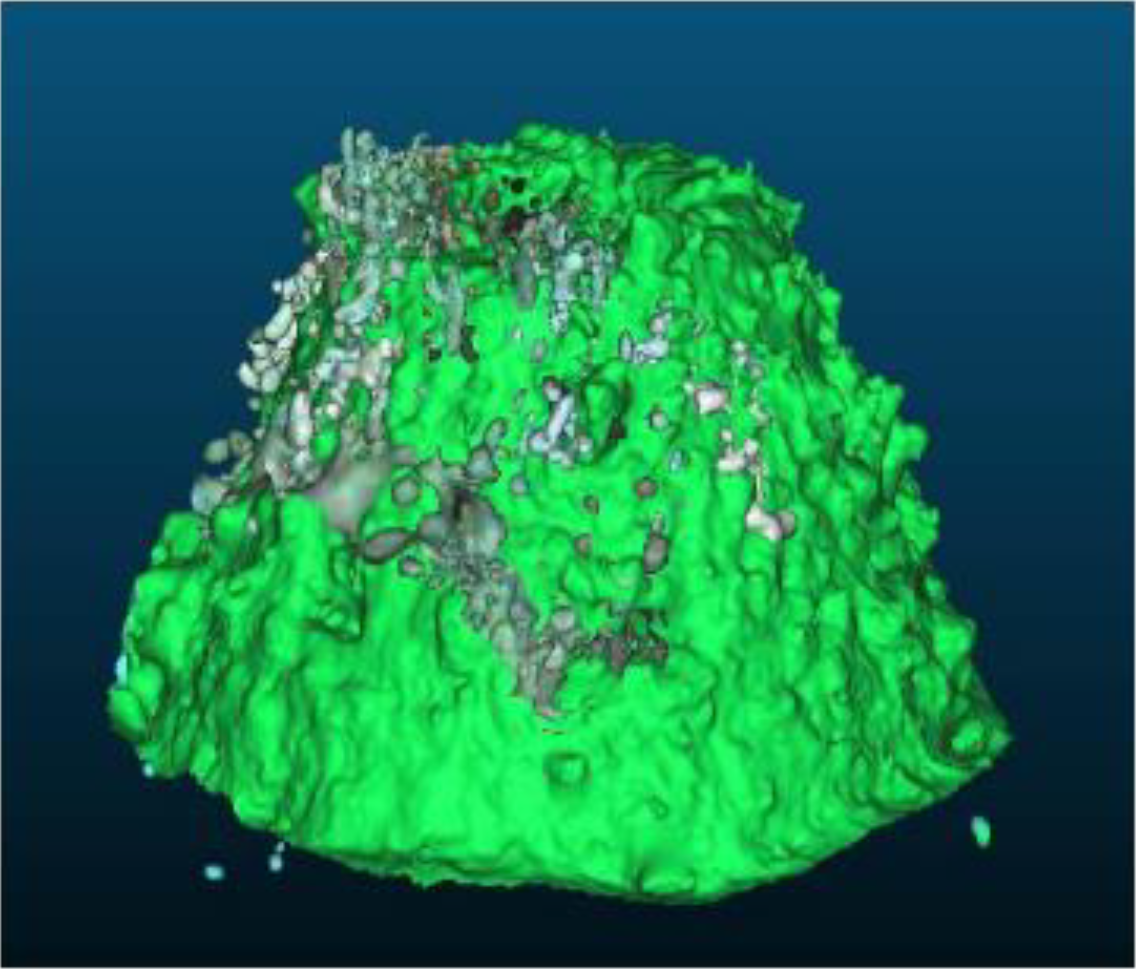
Example of the original model (light green) at 1m from Canon G7X Mark II overlaid with CLAHE model (dark green) at 1m. The CLAHE model can be seen as “noisy” due to the mesh including not just the structure itself.

**Figure S2.**
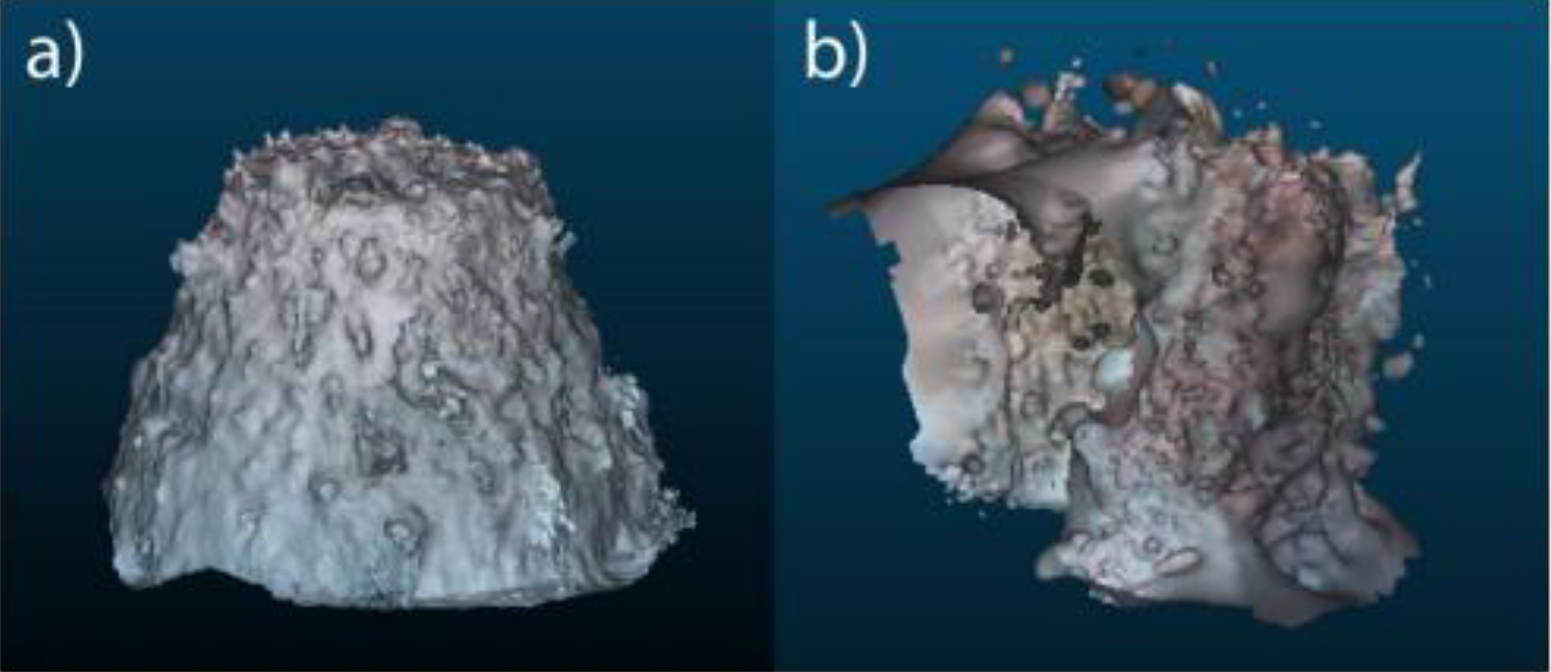
Models from combined Heights for 1m and 2m for a. GoPro Hero 8 original images. b. GoPro Hero 8 Histogram images.

